# Transcription initiation RNAs are associated with chromatin activation mark H3K4me3

**DOI:** 10.1101/265827

**Authors:** Matthew Hobbs, Christine Ender, Gregory J. Baillie, Joanna Crawford, Kelin Ru, Ryan J. Taft, Ryan J. Taft, John S. Mattick

## Abstract

Transcription initiation RNAs (tiRNAs) are small, predominantly 18 nt, RNAs whose biogenesis is associated with nucleosomes adjacent to active transcription initiation sites. These loci usually contain modified histones associated with transcription initiation, including histone H3 trimethylated at lysine 4 (H3K4me3). To further characterize the relationship of tiRNAs and H3K4me3 marked nucleosomes, H3K4me3-targeted RNA:chromatin immunoprecipitations were performed in a murine macrophage cell line, and small RNA sequence libraries were constructed and subjected to deep sequencing. The H3K4me3 libraries exhibited a distinct profile of read lengths with a noticeable enrichment of sequences 17-26 nt in length, with a peak at ∼18nt that included tiRNAs. These RNAs show clear enrichment of sequences that map to genomic features known to be associated with transcription initiation, including CAGE transcription initiation sites (TSSs), sites of RNAPII occupancy, and H3K4me3 sites. The distribution of sequences that map in the vicinity of TSSs is consistent with previous descriptions of tiRNAs; *viz*. a major peak at approximately 40 nt downstream of the TSS, and a minor broader peak approximately 150-200 nt upstream of, and on the opposite strand to, the TSS. These results show that tiRNAs are physically associated with H3K4me3-marked chromatin. tiRNAs may be markers of RNAPII pausing and it remains a possibility that their association with H3K4me3 is part of an epigenetic signaling system.

## Introduction

Although only about 2% of the mammalian genome is protein-coding, transcription is pervasive (Clark et al. 2011). Tens of thousands of individual non-protein-coding RNAs (ncRNA) transcripts have been catalogued, and a number of ncRNAs classes have been defined, including long intergenic RNAs (lincRNAs), and smaller species such as micro RNAs (miRNAs), small nucleolar RNA (snoRNAs), small interfering RNAs (siRNAs) and PIWI-associated RNAs (piRNAs).

We previously described two classes of small (predominantly ~18 nt) ncRNA, each associated with features of genes that are linked to nucleosome positioning. Splice site RNAs (spliRNAs) are derived from the 3? ends of exons adjacent to splice sites (Taft et al. 2010). Transcription initiation RNAs (tiRNAs) are associated with transcription start sites (TSSs), having a peak density at ~15 or ~35 nucleotides downstream of RNA polymerase II (RNAPII) transcription in humans and Drosophila respectively, consistent with their derivation from sequences immediately upstream of the first nucleosome in both species (Taft et al. 2009a; Taft et al. 2009c). tiRNAs are also commonly found near genomic CCCTC-binding factor (CTCF) binding sites in human and mouse (Taft et al. 2011), an architectural protein that helps define three-dimensional topological regions of the genome (Ong and Corces 2014). tiRNAs have also been identified at both processive and non-processive (e.g. enhancer) RNAPII loci (Taft et al. 2010), suggesting that their biogenesis is specifically tied to transcription initiation.

We have proposed (Taft et al. 2010) that tiRNAs are involved in marking or regulating the epigenetic landscape around transcription start sites, and speculated that since tiRNAs are derived from loci associated with transcription initiation they may be linked to chromatin mark histone H3 trimethylated on lysine 4 (H3K4me3) (Taft et al. 2010). Using RNA:chromatin immunoprecipitation and small RNA sequencing here we show tiRNAs are physically associated with H3K4me3 genomewide, consistent with a role as a potential epigenetic regulator.

## Results and Discussion

### Chromatin immunoprecipitation and generation of small RNAseq dataset

We performed H3K4me3-specific (K4) and control chromatin immunoprecipitations (ChIPs) in a mouse macrophage cell line, with or without exposure to lipopolysaccharide (LPS). From the immunoprecipitated chromatin we prepared small RNA sequence libraries, which were subjected to deep sequencing (Table 1).

**Table 1.** Short RNA sequencing libraries

| Designation | Antibody | LPS | Number of sequence reads |  |  |  |  |
| --- | --- | --- | --- | --- | --- | --- | --- |
|  |  |  | Raw | Clipped | NR<sup>1</sup> | NR mapped<sup>2</sup> | NR mapped uniquely<sup>3</sup> |
| in.2.- | none | no | 20,908,738 | 11,461,876 | 1,578,430 | 1,231,009 | 451,881 |
| in.2.+ | none | yes | 16,312,477 | 10,165,941 | 1,358,788 | 1,048,296 | 385,727 |
| IgG.- | IgG(rabbit) | no | 11,517,008 | 4,944,877 | 576,338 | 355,714 | 103,044 |
| IgG.+ | IgG(rabbit) | yes | 16,789,510 | 4,977,928 | 671,399 | 436,742 | 127,795 |
| IgG.M.- | IgG(mouse) | no | 15,585,036 | 6,789,798 | 855,751 | 566,892 | 169,289 |
| IgG.M.+ | IgG(mouse) | yes | 11,155,470 | 3,795,075 | 481,962 | 325,726 | 86,169 |
| K4.- | H3K4me3 | no | 12,093,363 | 8,319,574 | 1,413,629 | 1,134,944 | 572,931 |
| K4.+ | H3K4me3 | yes | 9,418,881 | 5,889,036 | 1,064,797 | 826,445 | 393,785 |

1. NR: Non-redundant clipped sequence reads
2. Disallowing mismatches, and allowing mapping to up to 100 genomic positions.

The distribution of lengths of non-redundant sequences (tags), which range from 16 nt to 76 nt (Figure 1). In all libraries there is a preponderance of lengths < ~30 nt with a modal length of 18 nt. The two K4 libraries, however, have a length distribution that is distinct from controls with enrichment of 17-26 nt sequences and depletion of sequences 28-58 nt in length. This difference is consistent with the K4 libraries being enriched in tiRNAs, which have a modal length of ~18 nt (Taft et al. 2009c).

**Figure 1.**
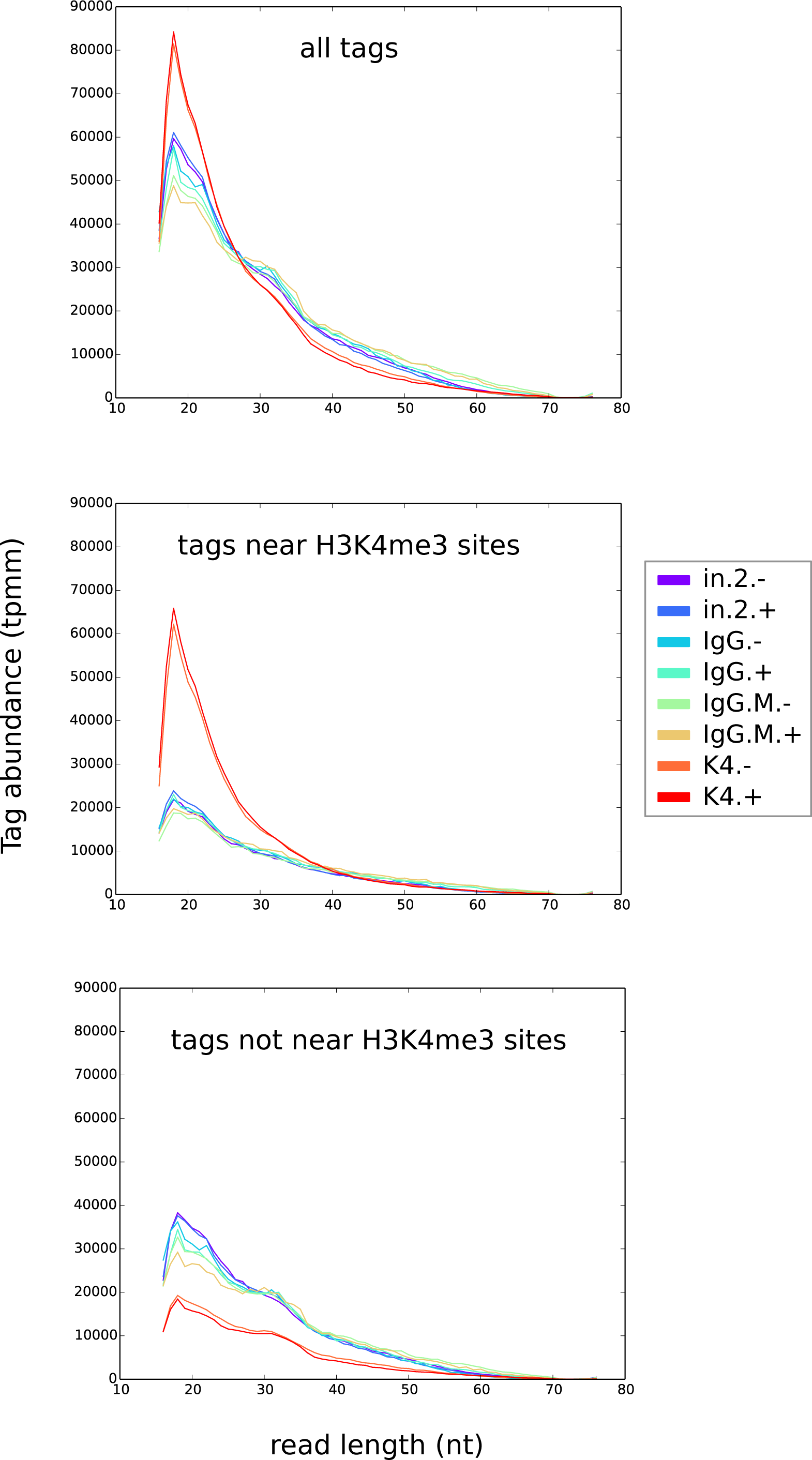
Distribution of lengths of small RNA sequences. Sequence abundance, measured as unique reads (tags) per million tags mapped (tpmm), is shown for all tags, as well as for tags partitioned according to their proximity to regions of peak occupancy by modified histone H3K4me3. The H3K4me3 occupancy locations were estimated using ChIP-seq data obtained from ENCODE.

### Small RNA proximity to sites of H3K4me3 occupancy

Small RNA sequencing reads were mapped to the mouse genome (mm10) and assessed for enrichment in the vicinity of H3K4me3 occupancy sites, as defined by ENCODE broad peak analyses of multi-tissue ChIP-seq experiments. The small RNAs from the K4 libraries are enriched (relative to controls) at these sites, but not at the occupancy sites of three other modified histones (H3K9me3, H3K27me3 and H3K36me3) examined (Figure 2 A). This enrichment is apparent in both the LPS+ and LPS- K4 libraries (Figure 1), and is consistent with our expectation that the two K4 libraries targeted RNAs that are physically associated with H3K4me3 sites within chromatin. Indeed, nearly half of the tags in the K4 libraries map near H3K4me3 sites (Figure 1). Furthermore, the enrichment at H3K4me3 sites is due to tags that are small in size (mode = 18 nt) (Figure 1) which is consistent with the previously observed length profile of tiRNAs, suggesting that tiRNAs are H3K4me3-associated sequences that are enriched in the K4 libraries.

**Figure 2.**
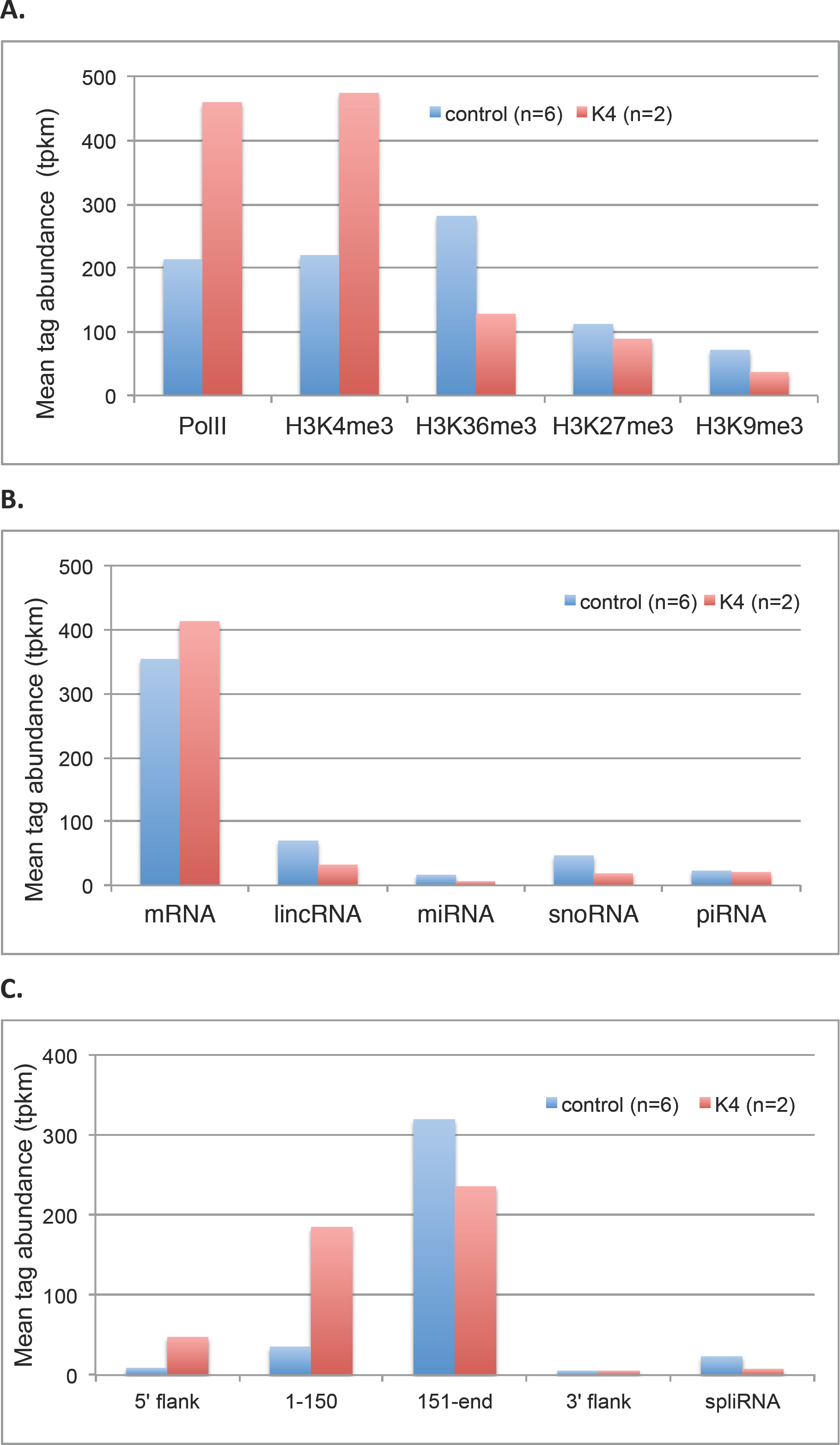
Abundance (measured as tags per thousand tags mapped (tpkm)) of tags which are located in the vicinity of various features within the genome. K4: mean tpkm from the two H3K4me3 specific libraries (K4.- and K4.+); control: mean tpkm from the remaining 6 non-H3K4me3-specific libraries. A. Tags mapping within 50bp of DNA occupancy regions of RNA polymerase II (PolII) and of four modified histones associated with different transcriptional states. The occupancy locations were estimated using ChIP-seq data obtained from ENCODE. B. Tags mapping within genes from 5 different classes of RNA. For the mRNA class, only tags mapping within exons of protein-coding genes are included, while for other RNA classes the whole gene is considered. C. A more detailed view of tags mapping in the vicinity of protein-coding genes. Tags were classified according to their occurrence in different parts of the gene: 5’ flank: 500 bp upstream flanking sequence; 1-150: the first 150 bp of the gene (but excluding any introns); 151-end: the remainder of the gene (again excluding introns); 3’ flank: 500 bp downstream flanking sequence. Also shown are spliRNAs, tags whose 3’end corresponds with the 3’ end of an exon of a protein-coding gene and whose transcriptional orientation is concordant (Taft et al. 2010).

### Mapping of small RNA sequences to genes encoding different classes of RNA

We compared the small RNA sequences with predicted transcript sequences from genome annotations covering several classes of RNA. Most unambiguously mapped tags were assigned to mRNA, although other classes of RNA, including lincRNA, snoRNA, piRNA and miRNA, are also represented in our dataset (Figure 2B). Also present in the dataset (not shown) are low levels of spliRNA-like sequences derived from the extremities of introns (Valen et al. 2011).

The K4 libraries are enriched in tags mapping to mRNA sequences, and relatively depleted in other classes (Figure 2B), and this enrichment is pronounced at the 5’ end of mRNA, and extends into the region upstream of protein-coding genes (Figure 2C). In contrast the K4 library is relative depleted of tags mapping to the residual part of mRNAs, and there is no enrichment downstream of protein-coding genes.

### Small RNA proximity to sites of transcription initiation

We looked at the position of small RNA sequencing reads relative to transcription start sites (TSSs) considering only TSSs of “sharp” promoters; that is, promoter regions where there is a single predominant TSS rather than a broad cluster of TSSs (Sandelin et al. 2007). The distribution of tag 3’ end positions relative to representative TSSs from FANTOM cap analysis gene expression (CAGE) data show that in both control and K4 libraries there is a peak of read abundance at approximately 40 nt downstream of the TSS, with read modal length 18 nt (Figure 3). This peak is more pronounced in the K4 libraries than in controls. The K4 libraries also show a second broader minor peak, located approximately 150-200 nt upstream of the TSS, also with 18 nt modal tag length, but comprising tags oriented in the opposite direction to TSS. This bimodal pattern is consistent with previous descriptions of tiRNAs (Taft et al. 2009a; Taft et al. 2009c; Valen et al. 2011), and is a global feature not restricted to TSSs associated with a particular class of gene (protein-coding, or non-coding) (results not shown).

**Figure 3.**
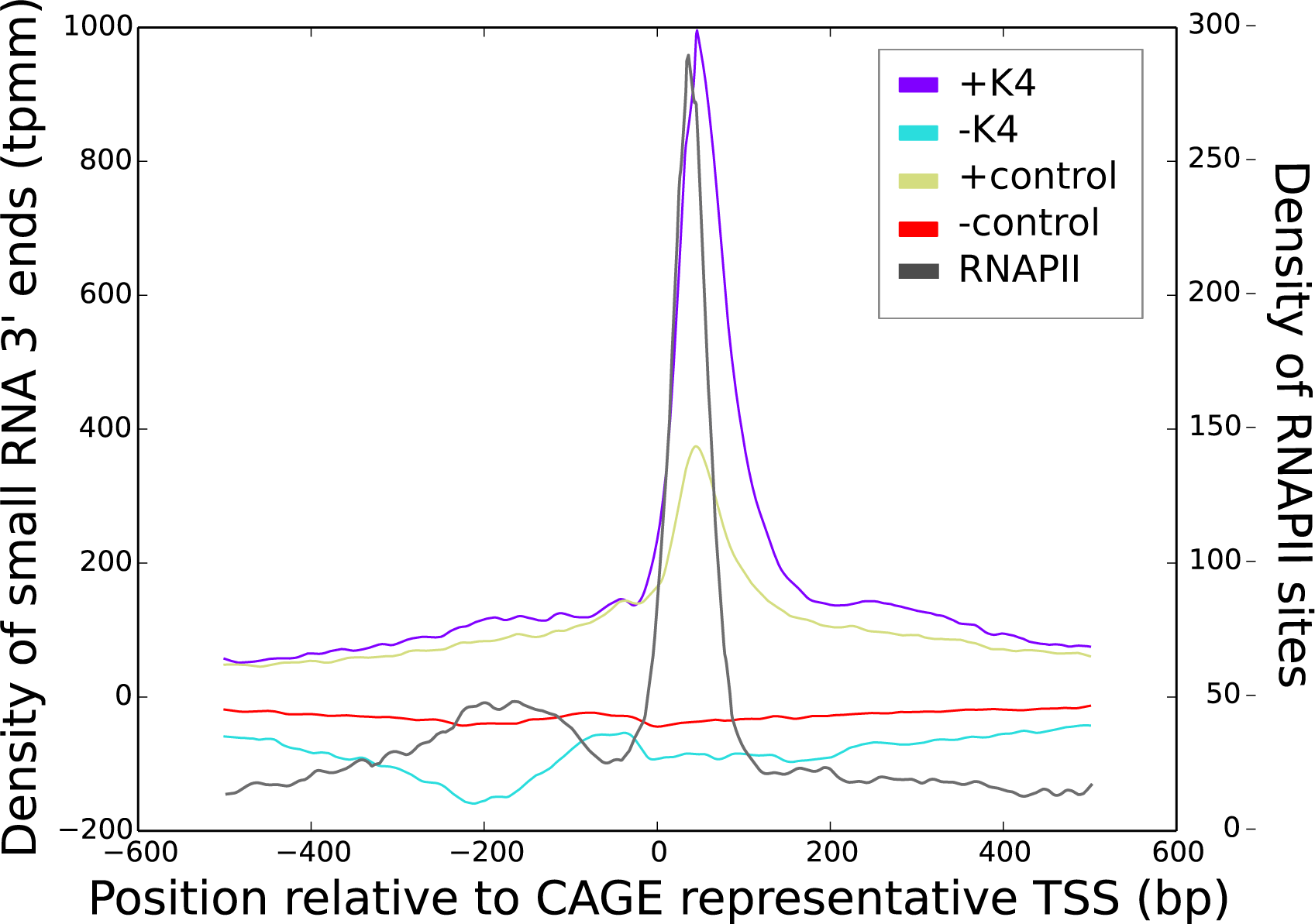
Abundance of mapped features in the vicinity of CAGE representative transcription start site (TSS). The CAGE data is limited to sharp promoters. The arrow shows the direction of transcription. Mean tag abundance (tpmm) in 6 control and 2 K4 libraries plotted against position of tag 3’ end. Values above the x-axis are from reads whose orientation is the same as that of the TSS, while those below are from reads with a discordant orientation. The major peak modal position is TSS+41. RNAPII site density is also plotted against site position, with the major peak modal position at TSS+31.

Taken together our results show that in comparison with controls the K4 libraries are enriched in tiRNAs. Furthermore this enrichment appears to be the major characteristic of the K4 libraries. Thus our ChIP sequencing provides evidence that tiRNAs are physically associated with H3K4me3-marked nucleosomes.

### tiRNA biogenesis and function

We have previously suggested that tiRNAs and spliRNAs are produced as a result of the propensity of RNAPII to backtrack and cleave when it encounters a nucleosome (Taft et al. 2009c). Others, however, have argued that tiRNAs may be the products of incomplete 5’ to 3’ exonucleolytic digestion given that the 3’ end of nascent RNA is protected by RNAPII (Gu et al. 1996; Li and Gilmour 2011; Valen et al. 2011), although we note that degradation products of other species of RNA may have biological significance (Taft et al. 2009b; Li et al. 2012).

Consistent with previous studies we see that there is a close relationship between the density of positions relative to the TSS of RNAPII and tiRNA sites (Figure 3B). It is possible that the occurrence of tiRNAs reflects RNAPII pausing, a process wherein transcription is reversibly halted after production a short (20-60 nt) nascent RNA. We previously considered this possibility but based on limited data available at that time concluded that the association of tiRNAs with RNAPII pausing was not strong (Taft et al. 2009a). Since then however numerous studies, including some employing techniques such as mNET-seq (Mayer et al. 2015) that greatly improve the sensitivity of detection of paused RNAPII, have shown that that RNAPII pausing is widespread (see reviews by (Kwak and Lis 2013; Jonkers and Lis 2015), and have also highlighted the importance of regulation of transcription at the level of elongation (Li and Gilmour 2011; Henriques et al. 2013; Scheidegger and Nechaev 2015). RNAPII pausing is commonly involved in transcription control of developmental and stimulus-responsive genes (Williams et al. 2015), and perhaps somewhat surprisingly is involved in processes such as exon skipping (Mayer et al. 2015) and alternative polyadenylation (Oktaba et al. 2015). Interestingly there is an association of CTCF with both tiRNAs (see above) and RNAPII pausing (Paredes et al. 2013).

We suggest the possibility that tiRNAs are sensors that report transcriptional roadblocks or checkpoints, and that they function within an epigenetic regulatory system. Henriques et al. (2013) have made a similar suggestion: that nascent transcripts associated with paused RNAPII might provide a target for regulatory processes such as chromatin remodeling or transcriptional elongation modulation.

### Conclusions

Our RNA sequencing provides evidence that tiRNAs are physically associated with chromatin marked with the transcription activation mark H3K4me3. tiRNAs may be markers of RNAPII pausing and as such could be candidate sensors of transcriptional elongation status in an epigenetic regulatory system.

## Materials and Methods

### Cell culture

Chromatin-associated small RNA preparations were prepared from the macrophage-like cell line RAW 264.7 (ATCC TIB-71; Raschke et al. (1978)). Cells were maintained on bacteriological plates in RPMI-1640 medium supplemented with 5% fetal bovine serum, 20 U/ml penicillin, 20 μg/ml streptomycin and 2 mM L-glutamine in a 37 °C incubator venting 5% CO_2_.

Prior to experiments, 8 Mio cells were plated on 15 cm tissue culture plates and incubated for 15 h. In the case of cells receiving LPS treatment, cells were stimulated 3 h prior to collection by addition of LPS (from Salmonella Minnesota, Sigma Aldrich) to a final concentration of 100 ng/ml.

### Cross-linking

Protein-polynucleotide complexes were cross-linked by treatment of cells with formaldehyde (1% v/v in PBS (137 mM NaCl, 2.7 mM KCl, 10 mM Na_2_HPO_4_, 2 mM KH_2_PO_4_ at pH 7.4)) for 4 min at room temperature. The formaldehyde solution was removed and residual formaldehyde quenched by incubation of cells with glycine (330 mM in PBS) for 5 min. The glycine solution was removed and the plates rinsed twice with PBS.

### Cell lysis and chromatin preparation

After fixation, cells were resuspended in isotonic buffer (20 mM HEPES pH 7.5, 250 mM sucrose, 3 mM MgCl2, 0.5 % NP40, 3 mM β-mercaptoethanol, protease inhibitor (Roche complete, EDTA free) and RNAsin) by gentle scraping, transferred to a Dounce homogenizer and disrupted with 12 strokes of the pestle (type B).

The lysate was pelleted by centrifugation for 7 min at 1,200 g at 4 °C. The pellet was resuspended in isotonic buffer, layered onto a sucrose cushion (880 mM sucrose, 0.5mM MgCl_2_) and centrifuged for 7 min at 2,000 g at 4 °C. The cushion was removed and the pellet resuspended in hypotonic buffer (10 mM HEPES pH 7.9, 10 mM KCl, 1.5 mM MgCl_2_, 0.5 mM DTT, 0.25 % v/v NP40, protease inhibitors (Roche) and RNAsin), transferred to 2 ml Eppendorf tubes and centrifuged for 7 min at 300 g at 4 °C. The pellets were resuspended in 250 μl glycerol storage buffer (10 mM Tris pH 7.5, 0.1 mM EDTA, 5 mM MgAc, 25 mM glycerol) and stored at -80 °C until required.

In order to obtain chromatin fragments of about 150 bp, nuclear preparations were diluted in digestion buffer (100 mM Tris pH 7.5, 50 mM KCl, 8 mM MgCl_2_, 3 mM CaCl_2_) and digested with micrococcal nuclease (MNAse, NEB) for 10 m at 37 °C with gentle agitation. The reaction was stopped by transfer to ice and addition of EDTA to a final concentration of 10 mM, and stored at -80 °C until required.

### Chromatin immunoprecipitation

MNase-digested material was sonicated (5 × 10 sec on, 10 sec off, using a bath sonicator). IP buffer (10 mM Tris pH 8.0, 150 mM NaCl, 1 % v/v Triton X-100, 0.5 % w/v SDS, 1 mM EDTA, protease inhibitor, SUPERaseIn RNase inhibitor (Life Technologies)) was added to give a total volume of 1,750 ml per tube, the lysate passed once through a 27” no dead volume needle and centrifuged in a table top centrifuge for 5 min at full speed, 4C °C to remove nuclear debris.

Immunoprecipitation was performed by incubation of lysate and control antibody (normal IgG) or H3K4me3-specific antibody (ChIPAb+ Trimethyl-Histone H3 (Lys4) ChIP validated rabbit monoclonal antibody, Merck Millipore catalog no. 17-614) overnight at 4 °C under gentle agitation. During the last three hours, superparamagnetic beads (Dynabeads protein G, Life Technologies), previously equilibrated in IP buffer, were added. After immunoprecipitation, beads were put through a series of washes, each wash conducted on ice for 5 min: binding buffer (50 mM HEPES pH 7.5, 20 mM EDTA, 0.5 % v/v Triton X-100, 25 mM MgCl2, 5 mM CaCl2, 50 units/ml SUPERase In), FA500 buffer (50 mM HEPES pH 7.5, 1 mM EDTA, 1 % v/v Triton X-100, 500 mM NaCl, 0.1 % Na deoxycholate, 50 units/ml SUPERase In), LiCl buffer (10 mM Tris pH 7.5, 1 mM EDTA, 1 % v/v Triton X-100, 250 mM LiCl, 0.5 % Na deoxycholate, 50 units/ml SUPERase In), Tes buffer (10 mM Tris pH 7.5, 1 mM EDTA, 250 mM NaCl, 50 units/ml SUPERase In), and finally PBS.

To remove protein the samples were digested with proteinase K (Thermo Scientific) in Proteinase K buffer (100mM Tris pH 7.5, 150 mM NaCl, 12.5mM EDTA, 1% SDS) at 65 °C for 1 h.

### RNA extraction

Nucleic acids were prepared by acidic phenol/chloroform extraction followed by ethanol precipitation, and a wash with 80 % v/v ethanol. To remove DNA samples were next digested with DNAse (Turbo DNase, Life Technologies) at 37 °C for 30 min. Samples were then again treated with acidic phenol/chloroform extraction followed by ethanol precipitation, and the pellets washed with 80 % v/v ethanol.

### RNA sequencing

Sequencing libraries were prepared using the TruSeq Small RNA protocol (Illumina). Sequences were generated using an Illumina NextSeq 500, with a 1×75bp run. Sequences in fastq format were submitted to the NCBI Short Read Archive and assigned run identifiers SRR2145009, SRR2145008, SRR2144988, SRR2144965, SRR2144964, SRR2144963, SRR2142112 and SRR2144533. Sequence reads were clipped and collapsed using programs from the FASTX toolkit v. 0.0.13 (http://hannonlab.cshl.edu/fastx_toolkit/index.html). Clipping was with fastx_clipper run with parameters “-a TGGAATTCTC -l 16 -i -Q33”. A set of non-redundant collapsed reads (tags) was then produced with fastx_collapser.

### Alignment of sequences to reference genome

Tag sequences were aligned to the GRCm38 (mm10) assembly of the mouse genome using bowtie2 (Langmead and Salzberg 2012) run with parameters “--local -k 100 -f -L 10 -N 1 -i L,1,0 -p 8 “. Alignments were converted to BAM format with samtools v. 0.1.18 (Li et al. 2009), sorted with picard v. 1.74 (http://broadinstitute.github.io/picard/), and then converted to BED and other formats with bedtools (Quinlan 2014).

### Proximity of tags to genomic features

Gene annotations were from the GENCODE project (Mudge and Harrow 2015). A comprehensive gene annotation file from release M4 of was downloaded from http://www.gencodegenes.org/mouse_releases/4.html.

Predicted mouse piRNA gene sequences were downloaded from piRNABank (Sai Lakshmi and Agrawal 2008) and used to create BED format files of piRNA gene positions.

CAGE data was from the FANTOM5 project (Lizio et al. 2015). A table of annotated CAGE peak positions for mouse samples was downloaded from http://fantom.gsc.riken.jp/5/datafiles/latest/extra/CAGE_peaks/mm9.cage_peak_phase1and2combined_coord.bed.gz. The representative transcription start positions given within this table were used in analyses.

Data from mouse ChIP-Seq experiments targeting POLR2A, H3K4me3, H3K9me3, H3K27me3 and H3K36me3 (ENCODE dataset accessions ENCSR000CFK, ENCSR000CFJ, ENCSR000CFF, ENCSR000CFD, ENCSR000CFE; generated in the Bing Ren lab, UCSD) were downloaded from the ENCODE Consortium portal website (Encode Project Consortium 2012; Sloan et al. 2016) as BED broadPeak format files which store regions of signal enrichment. Midpoints of the BED features were used in analyses.

Where positions of mouse genomic feature were given in mm9 coordinates they were converted to mm10 coordinates with liftOver (Hinrichs et al. 2006). Positions of aligned reads were compared with genomic features using programs from the bedops genome analysis toolkit (Neph et al. 2012). Features were visualized on the genome using the IGV genome browser (Thorvaldsdottir et al. 2013). Plots were produced with the Python 2D graphics package matplotlib (Hunter 2007).

## Acknowledgements

This research was supported under Australian Research Council’s *Discovery Projects* funding scheme (project number DP120103828). We are grateful to Kate Schroder for sharing cell lines and for advice on cell culture.

## References

Clark MB, Amaral PP, Schlesinger FJ, Dinger ME, Taft RJ, Rinn JL, Ponting CP, Stadler PF, Morris KV, Morillon A et al. 2011. The Reality of Pervasive Transcription. PLOS Biology 9: e1000625.

Encode Project Consortium. 2012. An integrated encyclopedia of DNA elements in the human genome. Nature 489: 57-74.

Gu W, Wind M, Reines D. 1996. Increased accommodation of nascent RNA in a product site on RNA polymerase II during arrest. Proceedings of the National Academy of Sciences of the United States of America 93: 6935-6940.

Henriques T, Gilchrist DA, Nechaev S, Bern M, Muse GW, Burkholder A, Fargo DC, Adelman K. 2013. Stable pausing by RNA polymerase II provides an opportunity to target and integrate regulatory signals. Molecular cell 52: 517-528.

Hinrichs AS, Karolchik D, Baertsch R, Barber GP, Bejerano G, Clawson H, Diekhans M, Furey TS, Harte RA, Hsu F et al. 2006. The UCSC Genome Browser Database: update 2006. Nucleic acids research 34: D590-598.

Hunter JD. 2007. Matplotlib: A 2D Graphics Environment. Computing in Science & Engineering 9: 90-95.

Jonkers I, Lis JT. 2015. Getting up to speed with transcription elongation by RNA polymerase II. Nature reviews Molecular cell biology 16: 167-177.

Kwak H, Lis JT. 2013. Control of transcriptional elongation. Annu Rev Genet 47: 483-508.

Langmead B, Salzberg SL. 2012. Fast gapped-read alignment with Bowtie 2. Nat Methods 9: 357-359.

Li H, Handsaker B, Wysoker A, Fennell T, Ruan J, Homer N, Marth G, Abecasis G, Durbin R, Genome Project Data Processing S. 2009. The Sequence Alignment/Map format and SAMtools. Bioinformatics 25: 2078-2079.

Li J, Gilmour DS. 2011. Promoter proximal pausing and the control of gene expression. Curr Opin Genet Dev 21: 231-235.

Li Z, Ender C, Meister G, Moore PS, Chang Y, John B. 2012. Extensive terminal and asymmetric processing of small RNAs from rRNAs, snoRNAs, snRNAs, and tRNAs. Nucleic acids research 40: 6787-6799.

Lizio M, Harshbarger J, Shimoji H, Severin J, Kasukawa T, Sahin S, Abugessaisa I, Fukuda S, Hori F, Ishikawa-Kato S et al. 2015. Gateways to the FANTOM5 promoter level mammalian expression atlas. Genome Biol 16: 22.

Mayer A, di Iulio J, Maleri S, Eser U, Vierstra J, Reynolds A, Sandstrom R, Stamatoyannopoulos JA, Churchman LS. 2015. Native elongating transcript sequencing reveals human transcriptional activity at nucleotide resolution. Cell 161: 541-554.

Mudge JM, Harrow J. 2015. Creating reference gene annotation for the mouse C57BL6/J genome assembly. Mamm Genome 26: 366-378.

Neph S, Kuehn MS, Reynolds AP, Haugen E, Thurman RE, Johnson AK, Rynes E, Maurano MT, Vierstra J, Thomas S et al. 2012. BEDOPS: high-performance genomic feature operations. Bioinformatics 28: 1919-1920.

Oktaba K, Zhang W, Lotz TS, Jun DJ, Lemke SB, Ng SP, Esposito E, Levine M, Hilgers V. 2015. ELAV links paused Pol II to alternative polyadenylation in the Drosophila nervous system. Molecular cell 57: 341-348.

Ong CT, Corces VG. 2014. CTCF: an architectural protein bridging genome topology and function. Nature reviews Genetics 15: 234-246.

Paredes SH, Melgar MF, Sethupathy P. 2013. Promoter-proximal CCCTC-factor binding is associated with an increase in the transcriptional pausing index. Bioinformatics 29: 1485-1487.

Quinlan AR. 2014. BEDTools: The Swiss-Army Tool for Genome Feature Analysis. Curr Protoc Bioinformatics 47: 11 12 11-11 12 34.

Raschke WC, Baird S, Ralph P, Nakoinz I. 1978. Functional macrophage cell lines transformed by Abelson leukemia virus. Cell 15: 261-267.

Sai Lakshmi S, Agrawal S. 2008. piRNABank: a web resource on classified and clustered Piwi-interacting RNAs. Nucleic acids research 36: D173-177.

Sandelin A, Carninci P, Lenhard B, Ponjavic J, Hayashizaki Y, Hume DA. 2007. Mammalian RNA polymerase II core promoters: insights from genomewide studies. Nature reviews Genetics 8: 424-436.

Scheidegger A, Nechaev S. 2015. RNA polymerase II pausing as a context-dependent reader of the genome. Biochem Cell Biol: 1-11.

Sloan CA, Chan ET, Davidson JM, Malladi VS, Strattan JS, Hitz BC, Gabdank I, Narayanan AK, Ho M, Lee BT et al. 2016. ENCODE data at the ENCODE portal. Nucleic acids research 44: D726-732.

Taft RJ, Glazov EA, Cloonan N, Simons C, Stephen S, Faulkner GJ, Lassmann T, Forrest AR, Grimmond SM, Schroder K et al. 2009a. Tiny RNAs associated with transcription start sites in animals. Nature genetics 41: 572-578.

Taft RJ, Glazov EA, Lassmann T, Hayashizaki Y, Carninci P, Mattick JS. 2009b. Small RNAs derived from snoRNAs. Rna 15: 1233-1240.

Taft RJ, Hawkins PG, Mattick JS, Morris KV. 2011. The relationship between transcription initiation RNAs and CCCTC-binding factor (CTCF) localization. Epigenetics & chromatin 4: 13.

Taft RJ, Kaplan CD, Simons C, Mattick JS. 2009c. Evolution, biogenesis and function of promoter-associated RNAs. Cell cycle 8: 2332-2338.

Taft RJ, Simons C, Nahkuri S, Oey H, Korbie DJ, Mercer TR, Holst J, Ritchie W, Wong JJ, Rasko JE et al. 2010. Nuclear-localized tiny RNAs are associated with transcription initiation and splice sites in metazoans. Nature structural & molecular biology 17: 1030-1034.

Thorvaldsdottir H, Robinson JT, Mesirov JP. 2013. Integrative Genomics Viewer (IGV): high-performance genomics data visualization and exploration. Brief Bioinform 14: 178-192.

Valen E, Preker P, Andersen PR, Zhao X, Chen Y, Ender C, Dueck A, Meister G, Sandelin A, Jensen TH. 2011. Biogenic mechanisms and utilization of small RNAs derived from human protein-coding genes. Nature structural & molecular biology 18: 1075-1082.

Williams LH, Fromm G, Gokey NG, Henriques T, Muse GW, Burkholder A, Fargo DC, Hu G, Adelman K. 2015. Pausing of RNA polymerase II regulates mammalian developmental potential through control of signaling networks. Molecular cell 58: 311-322.

